# Fractional Dynamics in Bioscience and Biomedicine and the Physics of Cancer

**DOI:** 10.1101/214197

**Authors:** Hosein Nasrolahpour

## Abstract

Almost all phenomena and structures in nature exhibit some degrees of fractionality or fractality. Fractional calculus and fractal theory are two interrelated concepts. In this article we study the memory effects in nature and particularly in biological structures. Based on this fact that natural way to incorporate memory effects in the modeling of various phenomena and dealing with complexities is using of fractional calculus, in this article we present different examples in various branch of science from cosmology to biology and we investigate this idea that are we able to describe all of such these phenomena using the well-know and powerful tool of fractional calculus. In particular we focus on fractional calculus approach as an effective tool for better understanding of physics of living systems and organism and especially physics of cancer.

## 1. Introduction

The concept of memory effect plays an important role in a large number of phenomena in different contexts and systems from biological structures to cosmological phenomena. For instance: memory effects in nanoscale systems [1, 2], optical memory effect [3], gravitational and cosmological memory effect (which means: the induction of a permanent change in the relative separation of the test particles when gravitational radiation passes through a configuration of test particles that make up a gravitational wave detector) [4], memory effects in gene regulatory networks [5], memory effects in economics [6] have been investigated. In addition other kinds of memory effect including: shape memory effect [7–10], magnetic shape memory effect [11] and temperature memory effect [12] have been also considered in recent years.

In all above mentioned references authors have used the standard calculus, however nowadays it is well-known that a natural way to incorporate memory effects in the modeling of various phenomena is using of fractional calculus. On the other hand almost all phenomena and structures in nature exhibit some degrees and levels of fractionality or fractality (low or high level or something between them), also nowadays it is well-known that there is a close relation between fractality and fractionality. In this work we investigate this idea that are we able to describe all of such these phenomena using the well-know and powerful tool of fractional calculus. Therefore for this purpose in the following, concepts of fractality and fractional dynamics are briefly reviewed respectively in Sec. 2.Then in Sec. 3 we introduce fractional calculus as a powerful tool for modeling of memory effects in different context and we present some important applications. At last, in Sec. 4, we will present some conclusions.

## 2. Fractality and fractionality

In the 1960s Mandelbrot introduced the concept of fractals. The main property of the fractal is non-integer Hausdorff dimension [13]. Fractality can exist either in space or in time. Fractals have many applications in various branches of science and engineering [14–17]. Nowadays, it is known that there is a close connection between fractals and fractional dynamics. Fractional dynamics is tied to the underlying fractal topology of space-time. Fractional dynamics studies the behavior of nonlinear physical systems that are [18–21]: out-of-equilibrium and described by differential and integral operators of non-integer orders. Equations containing fractal operators are used to analyze the behavior of systems characterized by: (1) power-law nonlinearity (2) power-law long-range spatial correlations or long-term memory (3) fractal or multi-fractal properties [22]. During the last years, the number of applications of fractional dynamics in science and particularly in physics has been steadily growing and include models of fractional relaxation and oscillation phenomena, anomalous transport in fluids and plasma, wave propagation in complex media, viscoelastic materials, non-Markovian evolution of quantum fields, networks of fractional oscillators and so on [23–29].

### 2-1. Fractality

A fractal is an object that has a non-integer dimension [13–15]. Fractals show self-similar structures. Such structures are introduced by using the concept of a reference structure and repeating itself over many scales. In general, the fractals structures are defined by an iterative process instead of an explicit mathematical formula [20]. As a simple example consider a closed (filled) unit equilateral triangle. Connect the mid-points of the three sides and remove the resulting inner triangle of size 0.5. By repeatedly removing equilateral triangles from the initial equilateral triangle, at the *n* th stage we will have an object consists of 3^*n*^ self-similar pieces (subsets) with magnification factor 2^*n*^. This is a well-known fractal that is called Sierpinski triangle or Sierpinski gasket. The fractal dimension of this fractal object is defined by

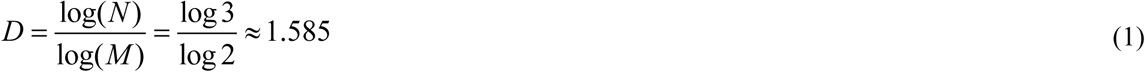

where *N* is the number of self-similar pieces (subsets) and *M* is the magnification factor. This number is a measure of irregularity and complexity of the fractal structure. Another simple case is the Cantor set which is a limiting set of points which results from discarding the middle third of each line segment in going from generation to generation, and starting from a line segment of unit length. The fractal dimension in the case of the Cantor set is

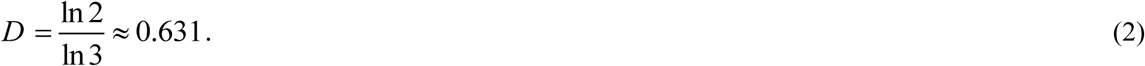

From classical mechanics to field theory and high energy physics fractality has many important applications [19–23]. For example fractal topology accounts naturally for breaking of discrete space-time symmetries and also it is able to account, at least in principle, for anomalous gauge charges, anomalous magnetic moment of massive leptons and enhanced cross-sections[21–23]. In recent years fractal geometry analysis has attracted attention of biologist as a useful and desirable tool to characterize the configuration and structure of biological structures. The considered issues include: fractal geometry analysis of configuration, structure and diffusion of proteins [24–29], fractal analysis of the DNA sequence, walks and aggregation [30–34] and fractal model of light scattering in biological tissue and cells [35].

### 2-2. Fractional dynamics

Fractional dynamics is a field in theoretical and mathematical physics, studying the behavior of objects and systems that are described by using integrations and differentiation of fractional orders, i.e., by methods of fractional calculus. Derivatives and integrals of non-integer orders are used to describe objects that can be characterized by: (1) a power-law non-locality (2) a power-law long-term memory (3) a fractal-type property [19]. As an example in the realm of classical physics we can consider the well-known diffusion phenomena. The most known diffusion processes is the normal diffusion. This process is characterized by a linear increase of the mean squared distance:

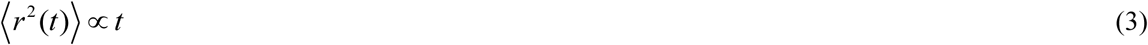

where *r* is the distance a particle has traveled in time *t* from its starting point. However there are many examples of phenomena in the natural sciences that violate this kind of behavior i.e. they are slower or faster than normal diffusion. In these cases (anomalous diffusions) the mean squared displacement is no longer linear in time:

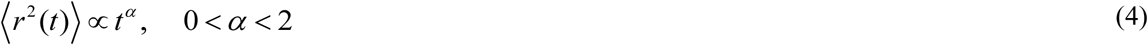

In recent years it is well known that generalization of the well-known diffusion equation and wave equation such that it includes derivatives of non-integer order with respect to time can describes phenomena that satisfy such a power law mean squared displacement. The fractional diffusion-wave equation[19] is the linear fractional differential equation obtained from the classical diffusion or wave equations by replacing the first-or second-order time derivatives by a fractional derivative (in the Caputo sense)[41–43] of order *α* with 0 < *α* < 2,

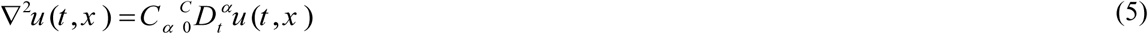

This equation describes diffusion-wave phenomena [36, 37] which is also called the anomalous diffusion such that we have the super-diffusion for 1 < *α* < 2, and sub-diffusion for 0 < *α* < 1. In above equations the fractional derivative of order *α*, *n-1* < *α* < *n*, *n* ∈ *N* is defined in the Caputo sense:

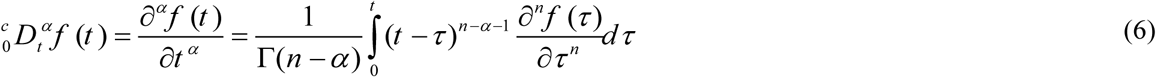

Where Γ denotes the Gamma function. For *α* = *n*, *n* ∈*N* the Caputo fractional derivative is defined as the standard derivative of order *n*. To derive a solution for a process described by an equation containing Caputo fractional derivatives, we need the initial conditions that can be written as:

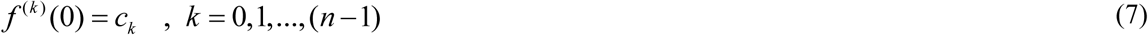

and because of this point that we are seeking the causal solution for natural phenomena we require that *f* (*t*) = 0 for *t* < 0.

Furthermore in the realm of high energy physics, it is generally believed that quantum field theory breaks down near the so-called Cohen-Kaplan threshold of ∝100 TeV as a result of exposure to large vacuum fluctuations and strong-gravitational effects. No convenient redefinition of observables is capable of turning off the dynamic contribution of these effects. For instance, it is known that the zero-point vacuum energy diverges quadratically in the presence of gravitation. Quantum field theory in Minkowski space-time discards the zero-point vacuum energy through the use of a normal time ordering procedure [44]. Because vacuum energy is gravitating and couples to all other field energies present at the quantum level, cancellation of the zero-point term is no longer possible when gravitational effects are significant. It is suggested that fractal geometry and fractional dynamics play a leading role in the description of TeV physics [22, 23]. As we see fractional dynamics and fractality are deeply tied to each other. In the following section we investigate the fractional calculus that is a major method in the consideration of fractional dynamical systems.

## 3. Fractional calculus in bioscience and biomedicine

As a physicist we always are able to model natural phenomena using systems of differential equations and nowadays it is well know that the advantage of fractional-order differential equation systems over ordinary differential equation systems is that they are more comprehensive and also incorporate memory effect in the model [38–44]. The kernel function of fractional derivative is called memory function [45]. Recently it is showed that the fractional model perfectly fits the test data of memory phenomena in different disciplines [46] they have found that a possible physical meaning of the fractional order is an index of memory. From this viewpoint fractional calculus has found many applications in new research on physics of biological structures and living organisms, from DNA dynamics [47–49] to protein folding [50], cancer cells [51], tumor-immune system [52], modeling of some human autoimmune diseases such as psoriasis [53], bioimpedance [54], spiking neurons [55], and also the transport of drugs across biological materials and human skin [56] and electrical impedance applied to human skin [57] and even modeling of HIV dynamics [58].

### 3-1. Fractional dynamics of cancer cells

Recently fractional order cancer model has been presented in Ref. [51]. In this work, they have studied the fractional-order model with two immune effectors interacting with the cancer cells. Without any doubt the behavior of most biological systems has memory and also some degrees and levels of complexity. As we mention in above sections the modeling of such these systems by fractional ordinary differential equations has more advantages than classical integer-order modeling, in which such effects are neglected. Studying immune system cancer interactions is an important topic (see [51 and references therein]). In Ref. [51] a new model for immune system has been proposed. In this model the authors have used two immune effectors and also they have considered cross reactivity of the immune system. The model with some modifications is as follows:

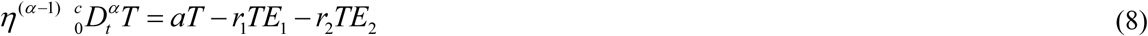

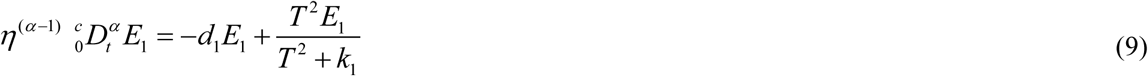

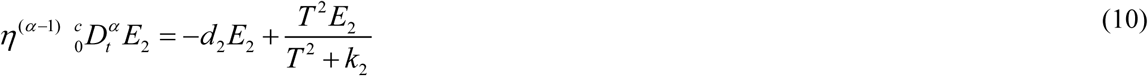

where *T* ≡ *T*(*t*) is the tumor cells, *E*_1_ ≡ *E*_1_(*t*), *E*_2_ ≡ *E*_2_(*t*) are the immune effectors, and *a*;*r*_1_;*r*_2_;*d*_1_;*d*_2_;*k*_1_;*k*_2_ are positive constants and *α* is the order of fractional derivative 0 < *α* < 1 and *η* is an arbitrary quantity with dimension of [second] to ensure that all quantities have correct dimensions. The interaction terms in the second and third equations of model (8-10) satisfy the cross reactivity property of the immune system. The equilibrium points of the system (8-10) are:

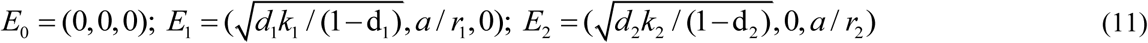

To avoid the non-biological interior solution where both immune effectors coexist, we assume that:

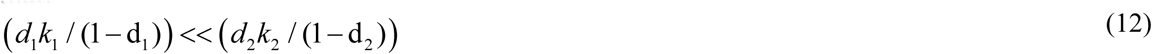

The first equilibrium *E*_0_ is the nave, the second *E*_1_ is the memory and the third *E*_2_ is endemic according to the value of the tumor size. Stability analysis shows that the nave state is unstable. However, the memory state is locally asymptotically stable if: d_1_< d_2_, and d_1_< 1 while the endemic state is locally asymptotically stable if: d_2_< d_1_, and d_2_< 1. Finally, based on the model described here the authors in Ref. [51] have showed that fractional order dynamical systems are more suitable to model the tumor-immune system interactions than their integer order counterpart. With this motivation in the next section we apply fractional calculus for the other area of new field of physics of cancer that is branching processes and cancer cells growth.

### 3-2 Cancer Growth

In this section we propose our new model for branching processes which are a class of simple models that have been used extensively to model growth dynamics of stem cells and cancer cells. Branching processes represent the most widely used approach to model population dynamics of cancer cells that can be defined in discrete or continuous time and with evolution rules that may or may not depend on time (for details see [60] and references therein). In this work we propose a new model for continuous time branching processes.

The natural generalization of the discrete time model for cells is the birth-and-death model where each cell has a probability *β*(*t*)*dt* of dying in the time interval (*t*, *t* + *dt*) and a probability *ϕ*(*t*)*dt* of dividing into two cells in the same time interval. Here the rates of cell death and division have been written as time dependent, but for simplicity we can assume they are constant.

The birth-and-death process can be used to compute the evolution of the size distribution of cancer cell colonies *p*(*s*, *t*), defined as the probability that a single cancer cell gives rise to a colony composed by *s* cells at time *t*, where time is measured in days. The probability density function *p*(*s*, *t*) evolves according to the following master equation:

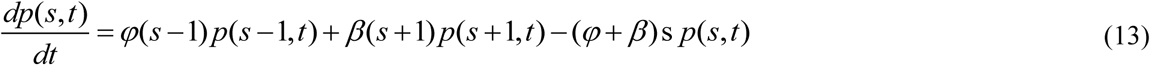

starting with an initial condition *p*(*s*,0)= *δ*_*s,1*_. From Eq. (13), we can obtain an equation for the average colony size 〈*s* 〉:

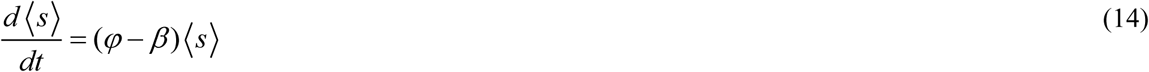

yielding an exponential growth:

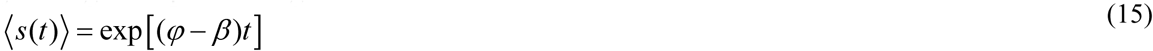

However branching processes is completely a complex biological phenomena and generally experimental dada deviates from the results of standard models and calculations so here we generalize Eq. (14) to the following fractional form:

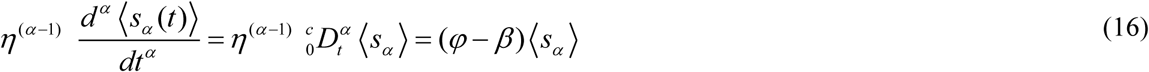

The solution of this equation can be written as:

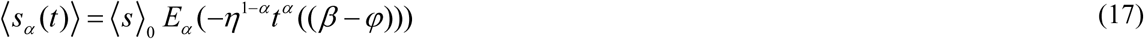

where

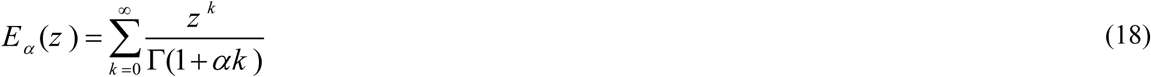

is one-parameter Mittag-Leffler function. For example for the case 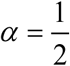 we have:

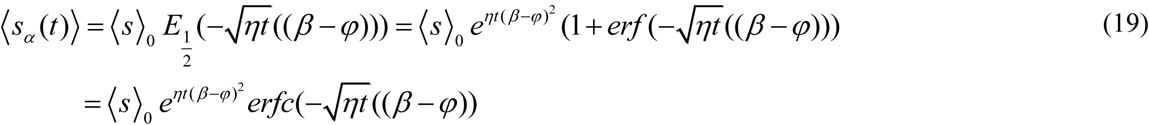

where *erfc* denotes the complimentary error function and the error function is defined as

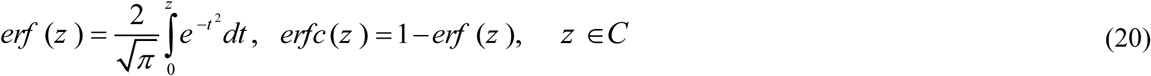

For large values of *z*,the complimentary error function can be approximated as

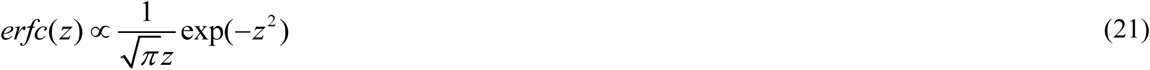

At asymptotically large times, and using Eq. (19) we have

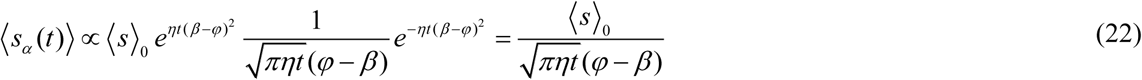

In many cases in nature and in particular in biological systems we may have low level of fractionality which mathematically means that the order of the fractional derivative *α* is close to a positive integer, namely, *α* = *n* − ε with small positive ε [59], then we will have:

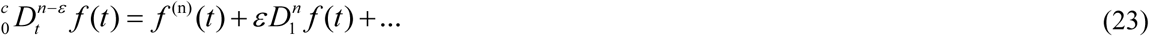

where 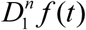 is:

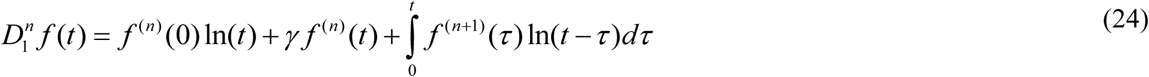

where *τ* < *t* and *γ* = 0.5552156… is the Euler constant.

In the case of low-level fractionality (i.e. the order of fractional derivative is *α* = 1 − ε) for our new model we will have:

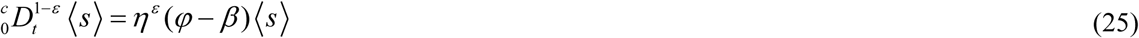

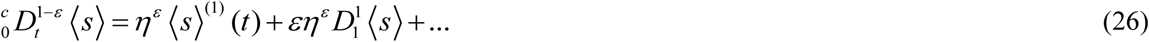

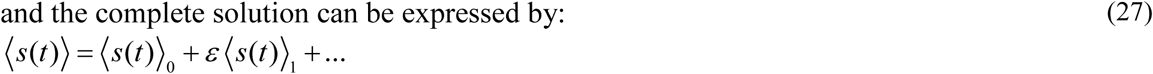

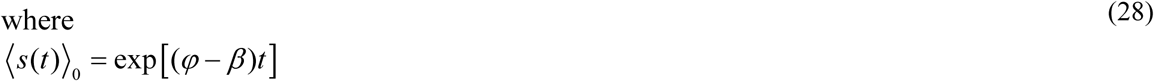

and for the correction term we have:

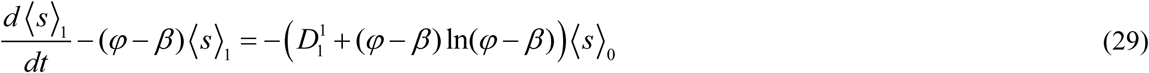

where,

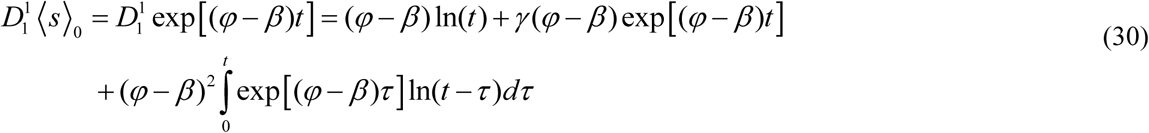

After Taking the Laplace transform of Eq. (29) we will have:

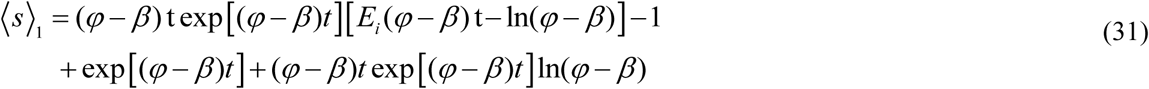

where *E*_*i*_(t) is the exponential integral defined by [61]:

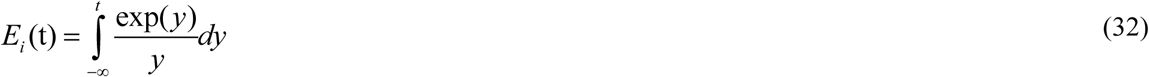

And has the following asymptotic form [61]:

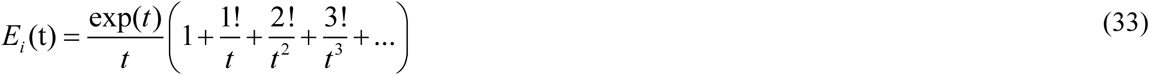

Therefore the complete solution can be expressed by:

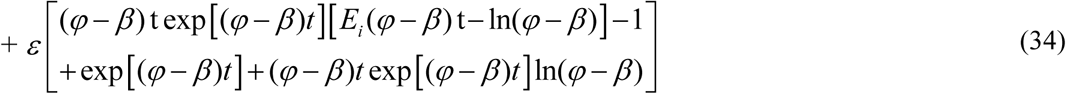

For small values of *t* we obtain:

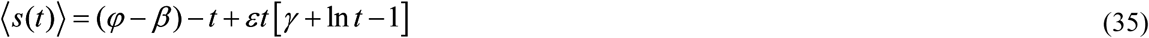

And for the asymptotic solution when *t* → ∞ we have:

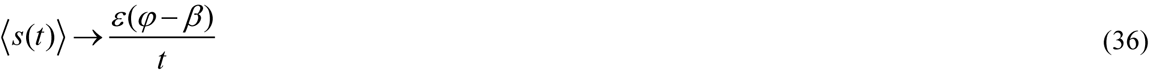

## 4. Conclusion

Fractals are measurable metric sets with a non-integer Hausdorff dimension. The main property of the fractal is non-integer Hausdorff dimension that should be observed on all scales. There is a close connection between fractals and fractional dynamics. Fractional dynamics is a field in physics and mechanics, studying the behavior of objects and systems that are described by using the fractional calculus. Fractals and fractional calculus generate parameters of arbitrary dimensions as well as arbitrary order of integration and differentiation. Therefore it is reasonable that we expect that behavior of the fractal objects and systems can be described by the fractional calculus approach. Almost all phenomena and structures in nature exhibit some degrees and levels of fractionality or fractality (low or high level or something between them) therefore it is reasonable to apply fractional calculus for them. As a scientist we always are able to model natural phenomena using systems of differential equations and nowadays it is well know that the advantage of fractional-order differential equation systems over ordinary differential equation systems is that they are more comprehensive and also incorporate memory effects in the models. In this work we have discussed about the fractional dynamics of cancer cells. We have proposed new model for the average colony size. We derived the average colony size in different conditions. We hope that these results and our future studies on the physics of biological structures and living systems help to better understanding of complexities that occur in such systems and organisms.

